# Delayed Reduction in Painful Sensations Evoked by Cervical tSCS Following High-Frequency Stimulation

**DOI:** 10.64898/2026.11.16.765970

**Authors:** Anna V. Makarova, Nadezhda Starodubtseva, Mikhail A. Lebedev

**Author notes:** Anna V. Makarova and Nadezhda Starodubtseva contributed equally to this work.

## Abstract

**Objective:** Transcutaneous spinal cord stimulation (tSCS) is a non-invasive neuromodulation technique with potential effects on pain perception. However, tSCS itself can evoke discomfort or pain at higher current intensities, and it remains unclear whether high-frequency tSCS alters the pain evoked by subsequent low-frequency stimulation. This study examined whether high-frequency cervical tSCS delivered at motor-threshold intensity is followed by changes in pain ratings elicited by low-frequency tSCS in healthy participants.

**Approach:** Twenty-five healthy participants completed a single experimental session. Individual sensory, motor, and pain thresholds were determined using single-pulse tSCS at the C6–C7 spinal level. Pain was assessed using 1-Hz stimulation delivered for 1 min at an individually defined intensity corresponding to 6–7 points on a 0–10 Numerical Rating Scale. Participants then received 80-Hz tSCS for 10 min at motor-threshold intensity, followed by three repeated pain-assessment sessions separated by 5-min intervals. Surface electromyography was recorded from the flexor carpi radialis and extensor carpi radialis. Pain analysis included 25 participants, whereas EMG analysis included 18 participants.

**Main results:** Pain ratings were significantly lower than baseline during the second and third post-stimulation assessments. In contrast, ratings obtained immediately after high-frequency stimulation did not differ significantly from baseline. EMG responses showed substantial inter-participant variability. No EMG metric showed a significant omnibus session effect after correction for multiple comparisons, although exploratory comparisons indicated transient increases in ECR AUC and mean rectified amplitude immediately after high-frequency stimulation.

**Significance:** These findings indicate that high-frequency cervical tSCS can exert a measurable short-term influence on pain perception. Pain ratings and stimulation-evoked EMG responses showed different temporal patterns and represent complementary, non-equivalent outcomes of spinal stimulation.

## 1 Introduction

Spinal cord stimulation (SCS) was introduced as a neuromodulation approach for pain control in the late 1960s and has since become an established therapeutic option for patients with refractory chronic pain, particularly neuropathic pain syndromes [1, 2]. Although epidural SCS remains the best-established approach to pain relief, its invasive implantation procedure and long-term hardware requirements have motivated growing interest in non-invasive spinal neuromodulation techniques [1, 3]. Transcutaneous spinal cord stimulation (tSCS) delivers current through surface electrodes placed over the spine and offers a non-surgical alternative to implanted epidural stimulation [4, 5, 6]. Neurophysiological and modeling studies indicate that tSCS can activate posterior-root afferents, which are spinal input structures similar to those recruited by epidural stimulation. Consequently, evoked motor responses are similar with both approaches [7, 8]. The physiological effects of tSCS depend strongly on stimulation site, electrode montage, pulse parameters, and outcome measures, and this variability has contributed to heterogeneous findings across studies [4, 5]. In particular, the influence of tSCS on pain perception and modulation of pain thresholds remains insufficiently explored.

Evidence from related forms of non-invasive spinal stimulation suggests that spinal neuromodulation can influence pain-related processing in both healthy participants and patient populations [1, 9]. In healthy participants, transcutaneous spinal direct current stimulation has been shown to inhibit nociceptive spinal pathway conduction, increase pain tolerance, and alter pain-related evoked responses, although results have not been consistent across experimental pain paradigms [10, 11, 12]. In patients with chronic pain, repeated anodal transcutaneous spinal direct current stimulation has been associated with reduced pain ratings and changes in neurophysiological markers of nociceptive processing, while preliminary evidence in people with limb amputation suggests that tSCS may reduce phantom limb pain and alter spinal excitability measures [9, 13, 14]. Together, these findings constitute a growing body of evidence that tSCS can influence pain processing, but the literature remains fragmented across stimulation modalities, patient groups, and outcome measures [1, 9, 15].

Pain modulation at the spinal level depends on interactions between nociceptive afferent input, segmental inhibitory and excitatory interneuronal processing, and descending supraspinal control, making the spinal cord a promising target for neuromodulatory interventions [16]. Spinal stimulation could therefore alter pain perception by changing dorsal horn processing, modifying sensory afferent excitability, and engaging inhibitory mechanisms that reduce nociceptive transmission [3, 17]. Because non-invasive spinal stimulation can also influence corticospinal and sensorimotor excitability, changes in pain thresholds may reflect both spinal and supraspinal mechanisms [18, 19].

EMG recordings are particularly informative in tSCS studies because they provide an objective readout of stimulus-evoked motor responses and can be used to estimate motor thresholds and quantify recruitment of spinal motor pathways [20, 21]. In cervical tSCS, stimulation primarily activates large-diameter posterior-root afferents and evokes muscle responses through transsynaptic recruitment of spinal motor pools rather than direct muscle stimulation [8, 22]. Thus, combining subjective pain ratings with EMG-based detection of motor responses makes it possible to dissociate sensory, motor, and pain-related effects of stimulation within the same individualized threshold-assessment protocol [22, 23].

Despite growing interest in non-invasive spinal stimulation, few studies have examined sensory perception, motor recruitment, and subjective pain ratings within the same individualized protocol, even though these outcomes are likely related but not interchangeable [9, 22]. Most existing work has focused either on motor rehabilitation and spinal excitability or on pain-related outcomes alone, making it difficult to determine how physiological recruitment thresholds relate to subjective pain experience during cervical tSCS [3, 4]. This gap is particularly relevant for healthy participants because stimulation can serve both as an experimental perturbation and as a model for probing the transition from sensation to motor recruitment and then to pain [12, 24]. Therefore, studies that compare sensory, motor, and pain thresholds and test whether high-frequency tSCS changes subsequent pain perception in healthy individuals are needed [9].

The present study addressed the discomfort induced by tSCS itself: as stimulation amplitude increases, participants report painful sensations. Consequently, tSCS intensity must be kept low to ensure participant comfort, which may prevent stimulation from reaching the intensity required to achieve the intended physiological effect. However, an analgesic effect may emerge during continued stimulation. To test this possibility, the present study examined whether tSCS produces after-effects that alter subsequent pain perception [17, 25]. We applied tSCS to healthy participants and combined subjective pain assessment with EMG-based detection of motor responses. Within an individualized stimulation framework comparing sensory, motor, and pain thresholds, we aimed to determine whether high-frequency cervical tSCS is followed by reduced pain ratings during subsequent low-frequency stimulation. This work may improve our understanding of the analgesic potential of tSCS and inform future research on its potential applications in clinical and experimental pain research [9, 22].

## 2 Materials and Methods

### 2.1 Participants

Twenty-five healthy participants (13 men and 12 women; mean age = 24.4 ± 4.0 years) participated in one experimental session lasting up to 60 min. The study was conducted in accordance with the Declaration of Helsinki and was approved by the Ethics Committee of the Institute for Information Transmission Problems of the Russian Academy of Sciences (protocol no. 2, 16 September 2025). All participants provided written informed consent, including consent for the use of anonymized data in research publications.

### 2.2 Experimental design

Electrical stimulation was administered using a NeoStim-5 stimulator (Cosyma, Russia), which delivered biphasic pulses with a total duration of approximately 2 ms. Two surface electrodes were used: the anode was positioned over the right clavicle, and the cathode was positioned over the C6–C7 spinal level. At each stimulation level, the following outcomes were recorded: (1) the presence of any sensation; (2) the presence of a visible motor response in the forearm muscles; and (3) subjective pain intensity, rated using a 0–10 Numerical Rating Scale (NRS).

To establish individual sensory, motor, and pain thresholds, single electrical pulses were delivered at 1 Hz with stepwise increases in current amplitude. Stimulation began at 1 mA and was increased in 1 mA increments up to 5 mA, followed by 5 mA increments thereafter. The sensory threshold was defined as the lowest current amplitude at which the participant reported any sensation, the motor threshold as the lowest amplitude that elicited a visible motor response, and the pain threshold as the lowest amplitude at which the participant reported any pain (NRS score greater than 0). Stimulation continued until participants reported distinctly painful sensations rated as 6–8 on the NRS, without exceeding a maximum current of 80 mA. To assess the reliability of subjective pain ratings, selected current amplitudes were subsequently presented again in a randomized order. Because current intensity was initially presented in ascending order, potential bias toward higher subjective ratings was considered during data interpretation.

The continuous stimulation sessions were presented in the following order:

1. Initial Pain Assessment Pain assessment was performed using the individually defined amplitude corresponding to a pain intensity of 6–7 on the NRS. Stimulation was delivered at a frequency of 1 Hz for 1 min. Participants were asked to rate their perceived pain intensity four times at equal intervals throughout the stimulation period.
2. High-Frequency Stimulation Session Following the initial pain stimulation, participants received high-frequency stimulation at 80 Hz. The stimulation amplitude corresponded to the motor threshold. The high-frequency stimulation protocol consisted of two phases:
  a. A gradual increase in amplitude from 50% to 100% of the target value over 1–3 min
  b. Continuous stimulation at the target amplitude for 10 min
3. Repeated Pain Assessment Sessions

Immediately after the high-frequency stimulation, participants underwent three additional pain assessment sessions. Each session used the same parameters as the initial pain stimulation (1 Hz, 1 min, amplitude corresponding to NRS 6–7). The sessions were separated by 5-min rest intervals.

### 2.3 Data acquisition and processing

Surface electromyography (sEMG) was used to record muscle activity from the flexor carpi radialis (FCR) and extensor carpi radialis (ECR). Two pairs of disposable Ag/AgCl ECG electrodes were placed over the muscle bellies following standard skin preparation. Signals were acquired in volts using an NVX-52 amplifier (Medical Computer Systems, Russia) at a sampling frequency of 50,000 Hz. No hardware filtering was applied other than a 50-Hz notch filter. Seven participants were excluded from the EMG analysis because poor signal quality prevented reliable identification of the stimulation artifact peak.

#### 2.3.1 EMG data processing

Raw EMG recordings were resampled to 20,000 Hz and standardized to two bipolar channels corresponding to the FCR and ECR. Stimulation onsets were identified using a combination of automated and manual procedures. Automated detection was based on the high-frequency components of the EMG signal associated with the stimulation artifact. Detection parameters were adjusted for individual recordings when necessary because signal characteristics varied across participants. Detected onsets were locally refined within a 1-ms window and visually inspected. Missed events were added manually, and spurious detections were removed before epoch extraction. EMG epochs were extracted from *−*25 to 50 ms relative to each detected stimulus onset. Baseline correction was performed using the interval from *−*25 to *−*1 ms. After baseline correction, the epochs were cropped to the interval from 2 to 50 ms. Before epoching, EMG signals were band-pass filtered from 20 to 500 Hz. Trial-level EMG metrics were calculated separately for each channel and epoch within a 2–25 ms response window. These included peak-to-peak amplitude (p2p), area under the rectified EMG curve (AUC), mean rectified amplitude, and response latency. Latency was defined as the first time point within the response window at which the signal exceeded a threshold of the baseline mean *±*3 standard deviations for at least 2 ms. The resulting trial-level dataset contained subject, session, epoch, channel, and metric values for all retained epochs. Because latency values contained a substantial proportion of missing observations, this metric was excluded from the main group-level EMG analysis.

#### 2.3.2 Statistical analysis

Statistical analysis was performed using a repeated-measures framework. To avoid pseudoreplication, individual EMG epochs were not treated as independent observations. Instead, trial-level EMG metrics were first aggregated for each participant, session, and channel. For each metric, median, mean, standard deviation, and count were computed, and the median value for each participant, session, and channel was used in the main analysis to reduce the influence of outliers. The primary EMG metrics included in the group-level analysis were peak-to-peak amplitude, area under the rectified EMG curve, and mean rectified amplitude. For each channel–metric combination, differences across the four repeated sessions were first assessed using the Friedman test. Only participants with complete data across all four sessions were included in each test. Resulting p-values were corrected for multiple comparisons using Holm correction. Exploratory pairwise comparisons between each post-stimulation session and session 0 were performed using Wilcoxon signed-rank tests. These comparisons were conducted independently of the omnibus Friedman test results. For each comparison, the number of participants, mean difference, standard deviation of differences, median difference, Wilcoxon statistic, and uncorrected p-value were obtained. P-values from pairwise tests were corrected both globally across all EMG pairwise tests and separately within each metric. The globally Holm-adjusted p-values were used for interpretation. Pain scores were analyzed separately at the participant level. For each participant, pain ratings obtained during the ascending and randomized threshold searches were averaged pointwise. The minimum (*P*_min_) and maximum (*P*_max_) of these averaged reference ratings were then used to normalize each pain rating *P* as *P*_norm_ = (*P −P*_min_)*/*(*P*_max_*− P*_min_). Participants for whom *P*_max_ = *P*_min_ were excluded from this normalization. For each session, four measured ratings from the pain stimulation period were retained, and their median normalized value was used as the participant-level summary measure. Session effects on pain scores were examined using paired Wilcoxon signed-rank tests for five selected comparisons: sessions 1, 2, and 3 versus session 0, session 2 versus session 1, and session 3 versus session 2. These exploratory comparisons constituted one family, and their p-values were adjusted using the Holm method. Additional exploratory checks of distributional assumptions were performed with Shapiro–Wilk tests and graphical inspection of score distributions and session-wise differences.

## 3 Results

### 3.1 Pain score distributions and statistical approach

Normalized pain scores were first examined across the four pain stimulation sessions. The distributions of normalized pain scores were similar across sessions and showed no pronounced skewness or multimodality (Fig. S1). Shapiro–Wilk tests did not detect significant deviations from normality at the session level (all *p≥* 0.330). However, at least one distribution of paired session differences deviated from normality (Fig. S2). Therefore, paired Wilcoxon signed-rank tests were used for the pain score comparisons.

### 3.2 Pain scores decreased after high-frequency stimulation

Normalized pain scores decreased across repeated pain stimulation sessions following the high-frequency stimulation block (Fig. 1(a)). Group-level normalized pain scores were highest during the initial pain stimulation session (session 0). Session 1 showed only a modest reduction relative to session 0, whereas sessions 2 and 3 showed lower values.

**Figure 1:**
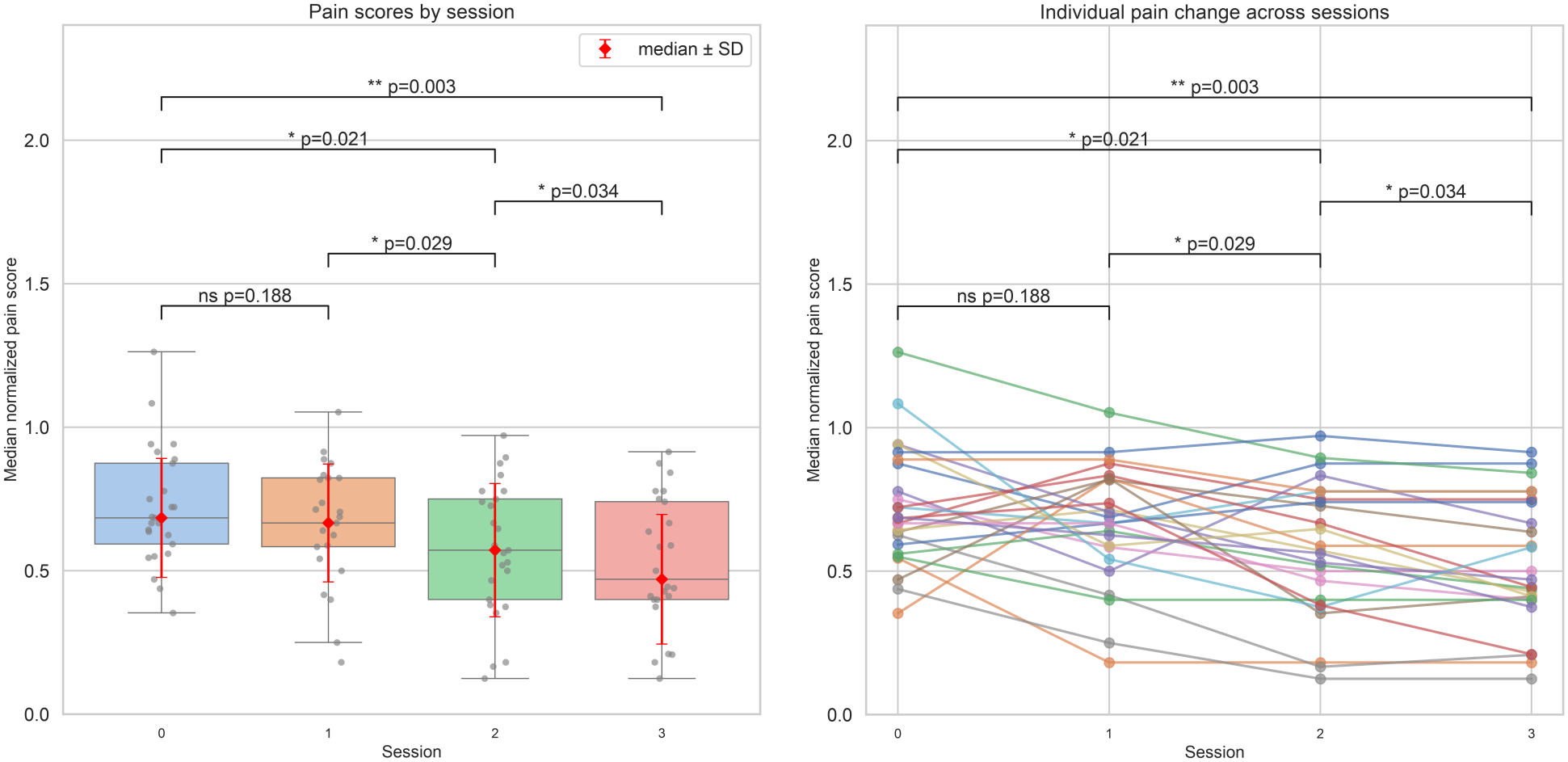
**(a)** Participant-level median normalized pain scores across the four experimental sessions. Boxplots show their distribution, gray dots indicate individual participants, and red diamonds indicate the group median *±*SD. Pain scores progressively decreased after the high-frequency stimulation block, with the largest reduction observed in the later post-stimulation sessions. **(b)** Individual participant trajectories of median normalized pain scores across sessions. Each line represents one participant. Although inter-participant variability was present, most participants showed stable or decreasing pain scores over time, with the strongest reduction typically observed in sessions 2 and 3. Brackets show Holm-adjusted p-values from paired Wilcoxon signed-rank tests.

Individual trajectories showed inter-participant variability but generally supported the group-level pattern, with most participants showing stable or decreasing pain scores across sessions (Fig. 1(b)). The largest reductions were observed in the later post-stimulation sessions. Pairwise comparisons indicated a delayed reduction in pain ratings (*n* = 25).Pain scores in session 1 did not differ significantly from session 0 (Wilcoxon *W* = 86.0, Holm-adjusted *p* = 0.188). In contrast, pain scores were lower in session 2 (*W* = 52.5, Holm-adjusted *p* = 0.021) and session 3 (*W* = 21.0,

Holm-adjusted *p* = 0.003) than in session 0. Exploratory comparisons between post-stimulation sessions showed lower pain scores in session 2 than in session 1 (*W* = 53.0, Holm-adjusted *p* = 0.029) and in session 3 than in session 2 (*W* = 18.0, Holm-adjusted *p* = 0.034). Scatterplots of individual values were consistent with these results. Values for session 1 remained close to the identity line relative to session 0 (Fig. 2(a)). In contrast, most values for sessions 2 and 3 fell below the identity line when compared with session 0, indicating lower normalized pain scores after high-frequency stimulation (Fig. 2(b, c)).

**Figure 2:**
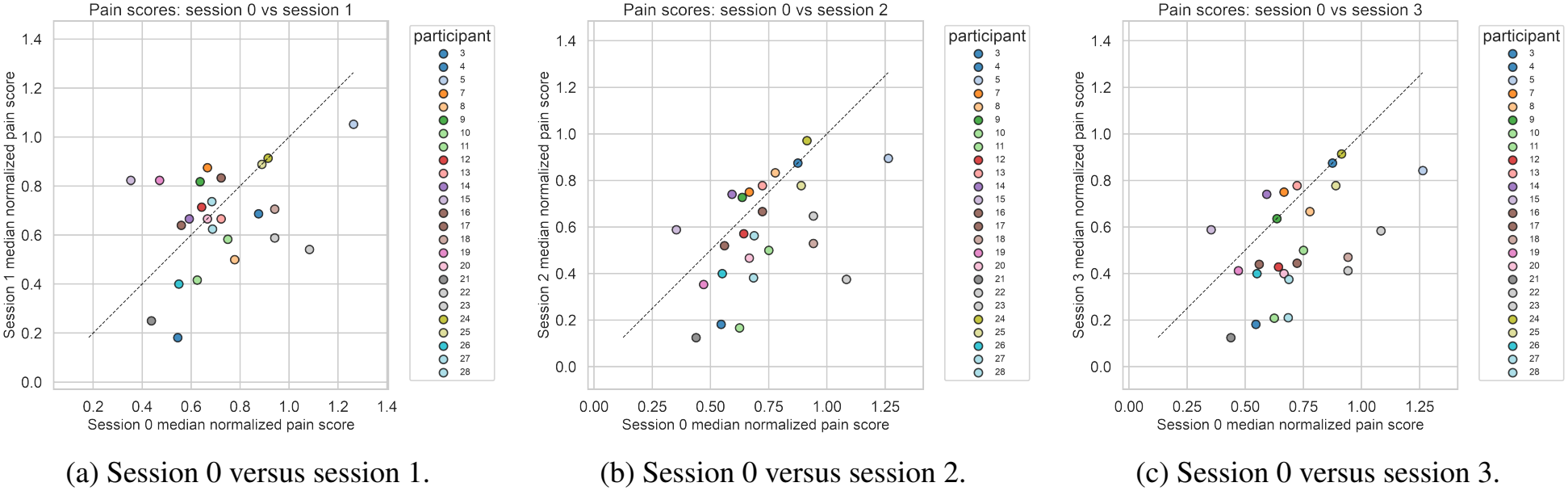
Scatterplots of participant-level median normalized pain scores at baseline and after high-frequency stimulation. Each point represents one participant, and the identity line is shown for reference. **(a)** Values in session 1 remained relatively close to the identity line, consistent with the absence of a strong immediate post-stimulation effect. **(b)** Most values in session 2 fell below the identity line, indicating lower pain scores than at baseline. **(c)** Most values in session 3 fell below the identity line, indicating the strongest reduction in the latest post-stimulation session.

### 3.3 EMG metrics showed high variability and limited session effects

EMG responses were analyzed using participant-level median values calculated for each session and channel. Three amplitude-based metrics were assessed: peak-to-peak amplitude, area under the rectified response curve, and mean rectified amplitude. Both ECR and FCR showed substantial inter-participant variability across sessions (Fig. 3(a–f)). In ECR, area under the rectified response curve and mean rectified amplitude showed a small increase from session 0 to later sessions, while peak-to-peak amplitude appeared higher in sessions 1 and 2 and closer to baseline in session 3 (Fig. 3(a–c)). However, these changes were small relative to the variability across participants. In FCR, the same metrics showed slightly higher values in sessions 1 and 2 compared with session 0, whereas session 3 remained close to or slightly above baseline depending on the metric (Fig. 3(d–f)). Distributions overlapped substantially across sessions. Friedman tests did not show significant session effects after Holm correction for any EMG metric (*n* = 18). For ECR, the Holm-adjusted p-values were 0.254 for peak-to-peak amplitude, 0.230 for AUC, and 0.203 for mean rectified amplitude; for FCR, the Holm-adjusted p-value was 0.176 for each metric. Exploratory baseline-versus-post comparisons showed higher ECR AUC in session 1 than in session 0 (Wilcoxon *W* = 18.0, globally Holm-adjusted *p* = 0.033) and higher ECR mean rectified amplitude in session 1 than in session 0 (*W* = 17.0, globally Holm-adjusted *p* = 0.028). No other pairwise comparison remained significant after global correction (all adjusted *p≥* 0.054). Thus, the EMG analysis provided limited evidence of an immediate increase in two ECR metrics but no consistent session-dependent modulation across muscles and metrics.

**Figure 3:**
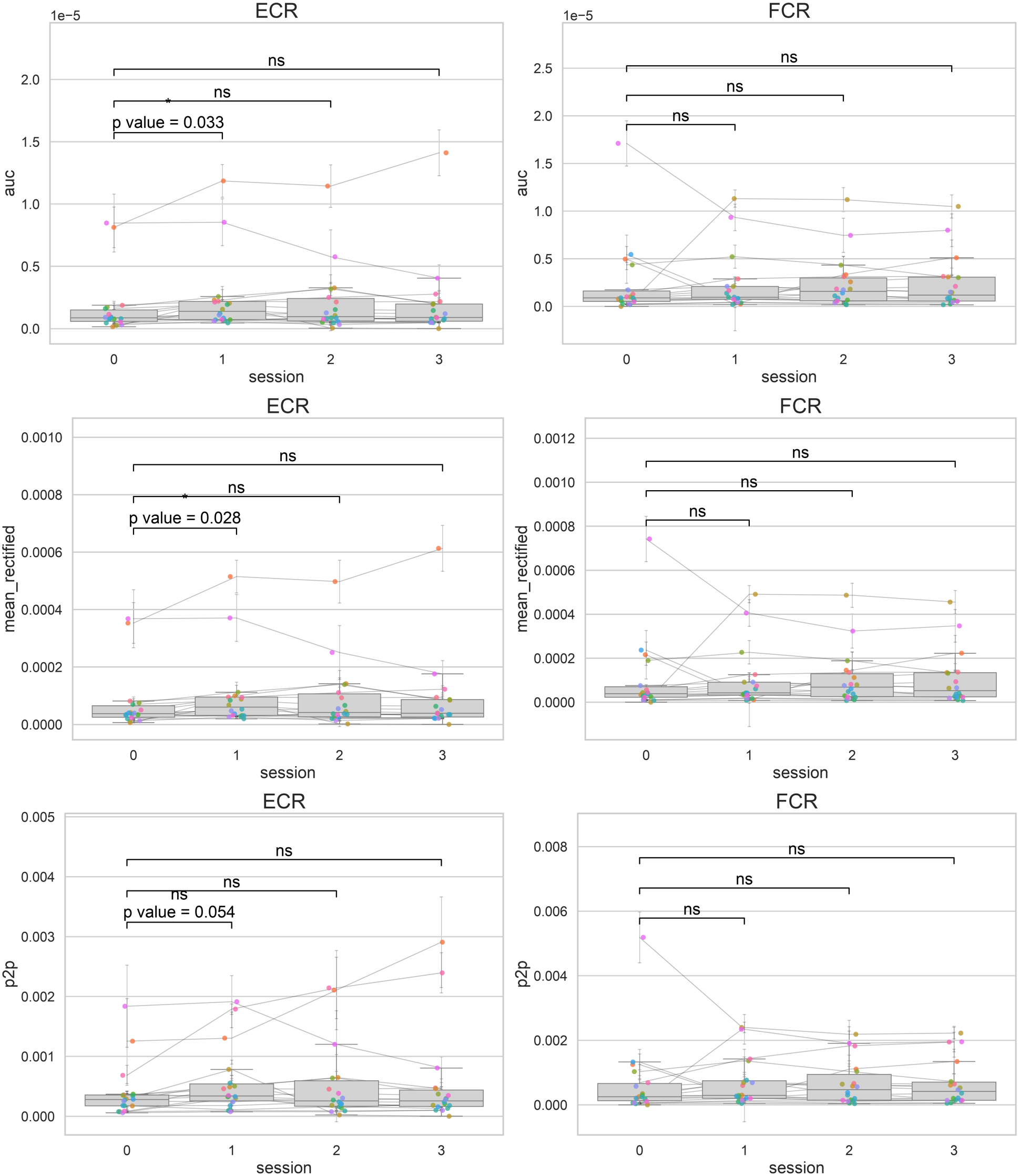
Session-wise distributions of participant-level median EMG metrics. Panels **(a–c)** show ECR AUC, mean rectified amplitude, and peak-to-peak amplitude (p2p), respectively; panels **(d–f)** show the corresponding FCR metrics. Colored dots represent individual participants, and lines connect measurements from the same participant across sessions. Brackets show globally Holm-adjusted p-values from exploratory paired Wilcoxon signed-rank tests comparing each post-stimulation session with session 0. Mild shifts in central tendency are visible across sessions, but the distributions are broad and highly variable.

### 3.4 Mean EMG waveforms showed heterogeneous responses

Averaged EMG waveforms showed stimulation-evoked components in both forearm muscles, but waveform morphology differed markedly between participants (Fig. S3).

Some participants showed clear short-latency biphasic or triphasic responses, whereas others showed broader, oscillatory, or early sharp components that may have included residual stimulation artifact. Within-participant difference waveforms did not show a consistent direction or morphology across sessions (Fig. S4).

## 4 Discussion

The present study examined pain perception and forearm motor responses within an individualized cervical tSCS protocol. The main finding was a delayed reduction in normalized pain ratings following high-frequency stimulation delivered at motor-threshold intensity. The reduction was not evident immediately after high-frequency stimulation but emerged during the later pain-assessment sessions. Omnibus tests did not show consistent session effects across EMG metrics. However, exploratory pairwise comparisons indicated transient increases in ECR AUC and mean rectified amplitude immediately after high-frequency stimulation. Thus, pain ratings and EMG metrics showed differing temporal and statistical patterns.

The delayed reduction in pain ratings is consistent with previous studies suggesting that non-invasive spinal stimulation may influence nociceptive processing both during stimulation and through short-lasting after-effects [16, 26, 27]. The temporal pattern supports the interpretation of a stimulation-related effect that developed after the intervention rather than an exclusively immediate perceptual response. Nevertheless, replication under sham-controlled or counterbalanced conditions will be important to assess the possible contributions of habituation and other time-dependent factors.

One possible explanation is that 80-Hz stimulation influenced nociceptive processing through spinal and/or supraspinal mechanisms [16, 26, 27]. Because stimulation was delivered at an individually defined level associated with a visible motor response, it may have recruited sensorimotor spinal circuits while remaining below the individually defined painful stimulation intensity [5, 6]. However, these mechanisms were not measured directly, and the present design cannot distinguish segmental spinal effects from broader supraspinal contributions.

The EMG findings provide limited evidence of an immediate, muscle-specific change rather than a uniform modulation of forearm motor responses. No EMG metric showed a significant omnibus session effect after correction, whereas exploratory comparisons identified increases in ECR AUC and mean rectified amplitude in session 1 relative to session 0. These changes were not consistently present in later sessions or in FCR. The physiological substrates of pain modulation and forearm motor recruitment overlap but are not identical, and the effects of tSCS depend on electrode montage, intensity, and anatomical variability [8, 22, 23, 28, 29]. Accordingly, pain ratings and motor responses provide complementary, non-equivalent readouts of the effects associated with spinal stimulation.

The physiological effects of the protocol were heterogeneous across participants. Some averaged EMG waveforms contained clear biphasic or triphasic short-latency components, whereas others were broader or more oscillatory, and early components in some recordings may have included residual stimulation artifact. The analysis emphasized stimulation-onset detection and a conservative response window rather than aggressive artifact removal. This strategy supported consistent epoch alignment but may have reduced sensitivity to subtle changes in early response morphology. Therefore, the absence of consistent omnibus EMG effects should not be interpreted as evidence that physiological modulation was absent [5, 30, 31, 32].

An important feature of the study was the comparison of sensory perception, visible motor responses, and painful stimulation levels within the same individualized protocol. The threshold search informed an initial pain-assessment session, a high-frequency motor-threshold stimulation phase, and repeated post-stimulation pain-assessment sessions. This design allowed us to examine whether stimulation calibrated according to an observable motor response was followed by changes in pain perception measured at a separate participant-specific pain level. Pain ratings showed a delayed reduction, whereas EMG findings were limited to transient exploratory changes in two ECR metrics, indicating that these measures capture related but non-equivalent aspects of the response to cervical tSCS [22, 23, 28, 29].

Several limitations should be acknowledged. The study involved healthy participants and experimentally induced pain produced by low-frequency cervical stimulation; the findings therefore cannot be directly generalized to chronic or neuropathic pain [26, 27]. The absence of a sham condition means that, although the temporal pattern is consistent with a stimulation-related effect, contributions from habituation, expectation, adaptation, or other time-dependent factors remain possible. The randomized threshold-search pass reduced potential ascending-order bias during individual calibration but did not control for effects arising across the four sequential pain-assessment sessions. Participant-specific normalization was necessary because pain ratings are subjective and vary substantially between individuals, although it also made the normalized values dependent on each participant’s threshold-search ratings. In addition, EMG analysis was limited to 18 of the 25 participants because seven recordings had insufficient signal quality, and latency was excluded from group-level analysis because of missing values. The sample size allowed detection of differences in pain ratings but may have provided limited sensitivity to small and heterogeneous EMG effects. Finally, the high-frequency and pain-inducing stimulation protocols differed in stimulation frequency and in the method used to select stimulation intensity, limiting mechanistic attribution.

Future studies should test the reproducibility of the delayed reduction in pain ratings using sham-controlled, counter-balanced, or crossover designs. Such studies could clarify the relative contributions of high-frequency stimulation, repeated exposure, and expectancy effects. It would also be valuable to examine alternative stimulation frequencies, durations, and montages, because tSCS effects depend strongly on stimulation parameters [22, 23, 28, 29]. Larger samples and improved EMG signal quality would permit more detailed analyses of waveform morphology, latency, recruitment curves, and participant-level response patterns.

Overall, the present findings support the idea that cervical tSCS can exert a measurable short-term influence on pain perception in healthy individuals, even when stimulation is calibrated according to motor rather than pain threshold. At the same time, the results indicate that pain ratings and stimulation-evoked forearm EMG responses should not be treated as equivalent physiological outcomes. Instead, pain ratings and motor responses provide complementary, non-equivalent readouts of the effects associated with spinal stimulation [16, 23, 26, 27, 29].

## 5 Conclusion

In this study, cervical tSCS was examined both as a stimulus capable of evoking pain and as a potential intervention for modulating subsequent pain perception in healthy participants. High-frequency tSCS delivered at motor-threshold intensity was followed by a delayed reduction in normalized pain ratings during subsequent low-frequency stimulation. This reduction was not evident immediately after high-frequency stimulation but emerged during the later post-stimulation assessments. EMG responses showed substantial inter-participant variability and no significant omnibus session effects. Exploratory comparisons nevertheless indicated transient increases in ECR AUC and mean rectified amplitude immediately after high-frequency stimulation. Thus, pain ratings and forearm motor responses showed different temporal and statistical patterns. Together, these findings support the short-term pain-modulating potential of non-invasive cervical tSCS and demonstrate the value of combining sensory, motor, pain, and EMG measures within an individualized experimental framework.

## 6 Author Contributions

Anna V. Makarova: Conceptualization, Methodology, Data curation, Formal analysis, Visualization, Writing–original draft. Nadezhda Starodubtseva: Conceptualization, Methodology, Data curation, Formal analysis, Writing–original draft. Mikhail A. Lebedev: Writing–review and editing, Supervision.

## 7 Competing Interests

The authors declare no competing interests.

## 8 Supplementary Materials

**Figure S1:**
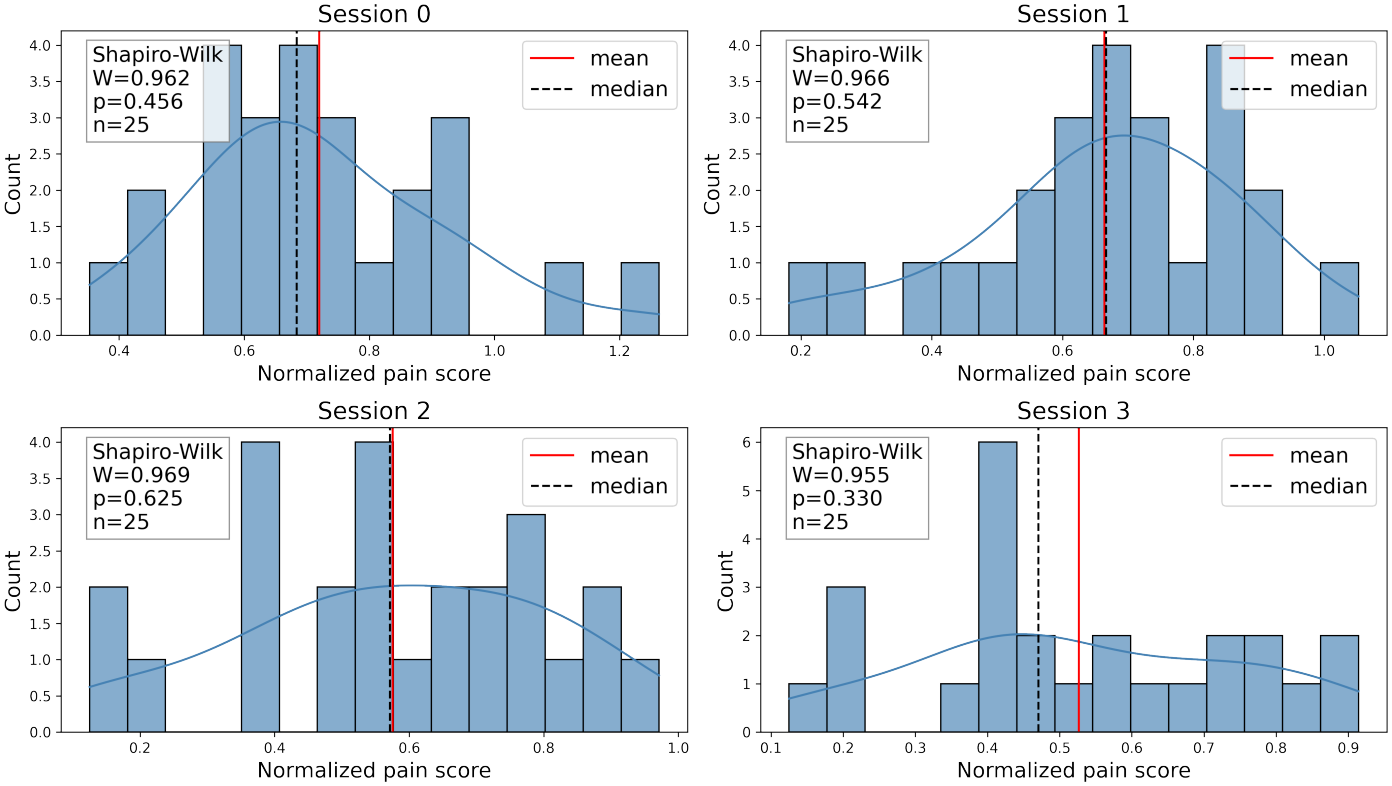
Distributions of participant-level median normalized pain scores in each session. Histograms and density curves are shown together with mean and median markers and Shapiro–Wilk test statistics. Shapiro–Wilk tests did not detect significant deviations from normality at the session level.

**Figure S2:**
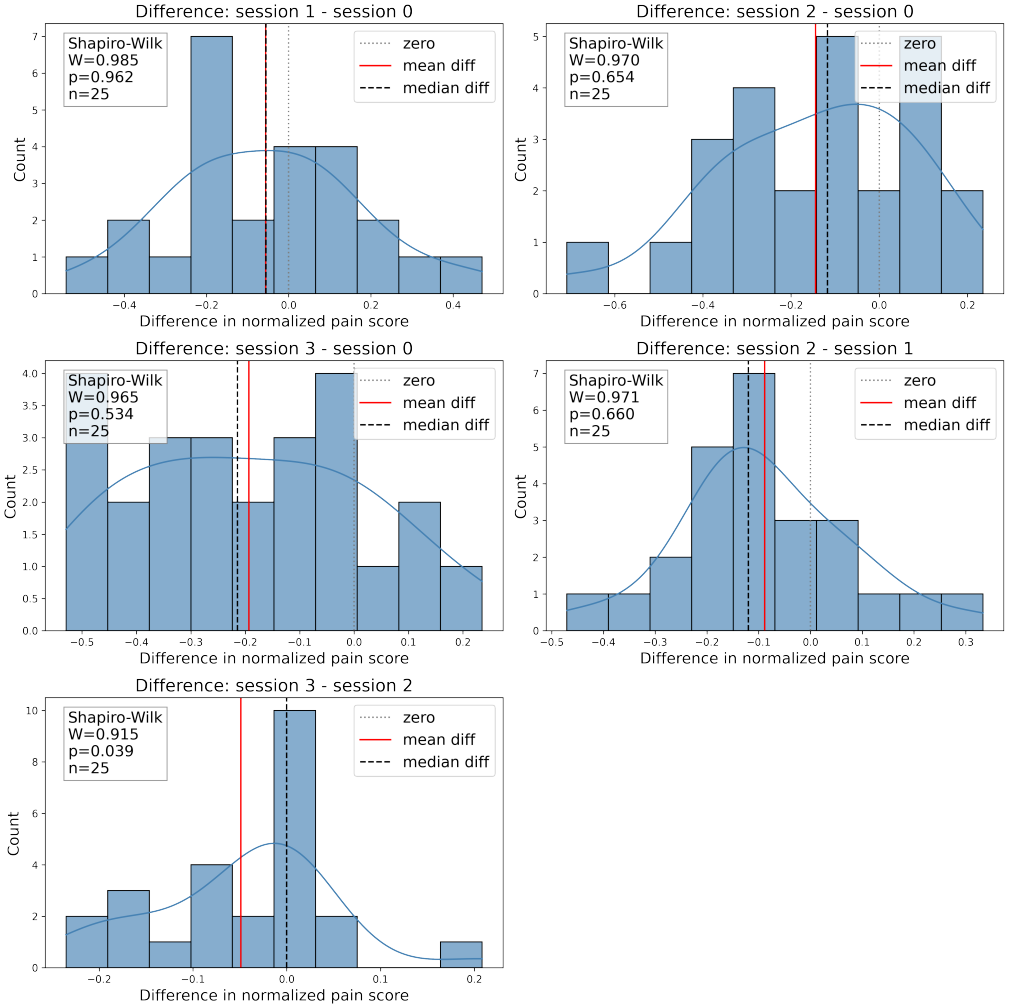
Distributions of pairwise differences in participant-level median normalized pain scores, calculated as the later session minus the earlier session. Histograms and density curves are shown together with mean and median difference markers, the zero reference line, and Shapiro–Wilk statistics. At least one comparison deviated from normality, supporting the use of non-parametric paired testing.

**Figure S3:**
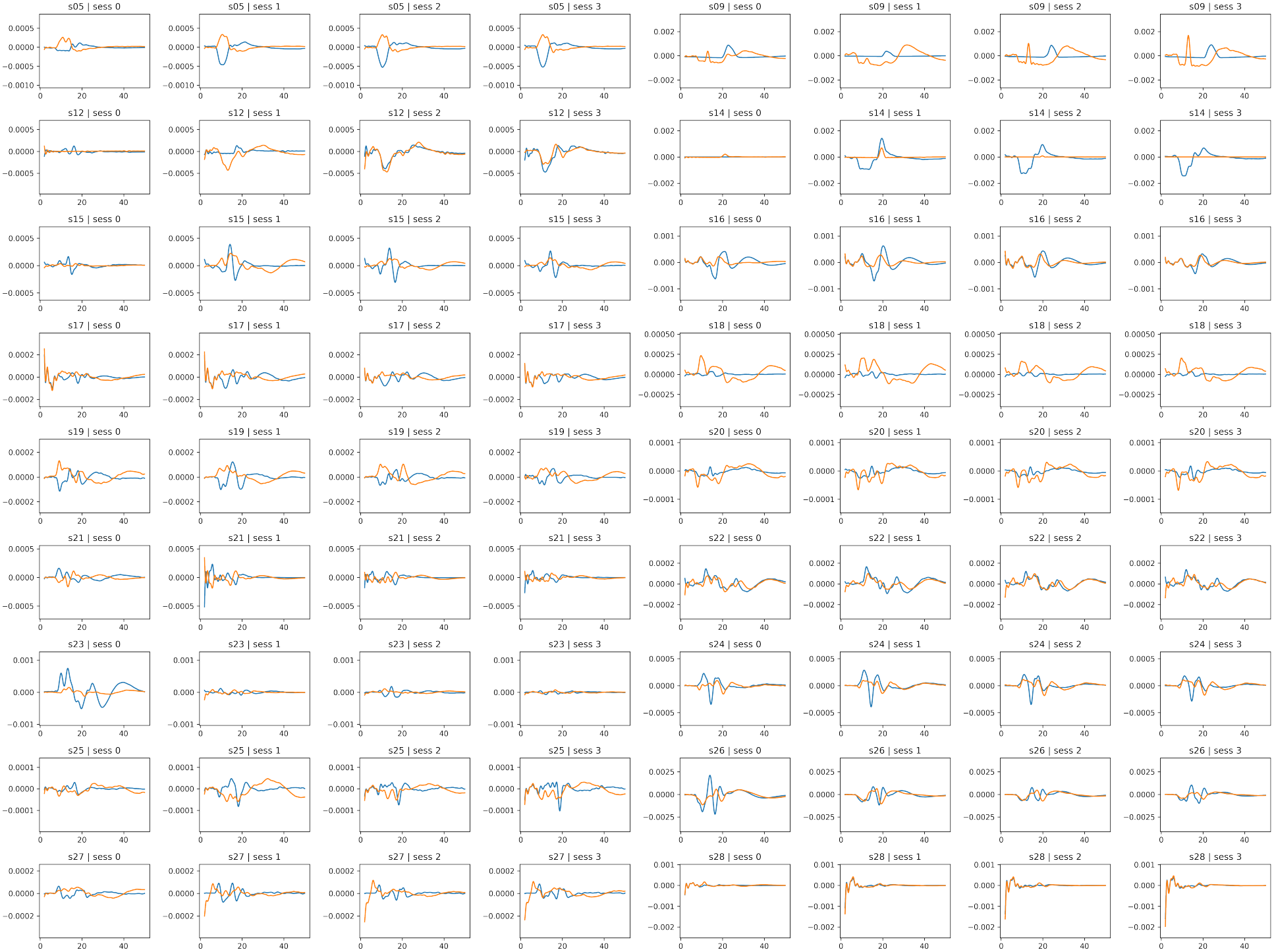
Trial-averaged stimulation-evoked EMG waveforms for individual participants and sessions. Each row corresponds to one participant, and the four adjacent panels show sessions 0–3. Blue traces represent FCR and orange traces represent ECR. The horizontal axis indicates time relative to stimulus onset (ms), and the vertical axis indicates EMG amplitude (V). Response morphology was generally more similar within than between participants but also varied across sessions, illustrating marked inter-participant heterogeneity in waveform shape and response magnitude.

**Figure S4:**
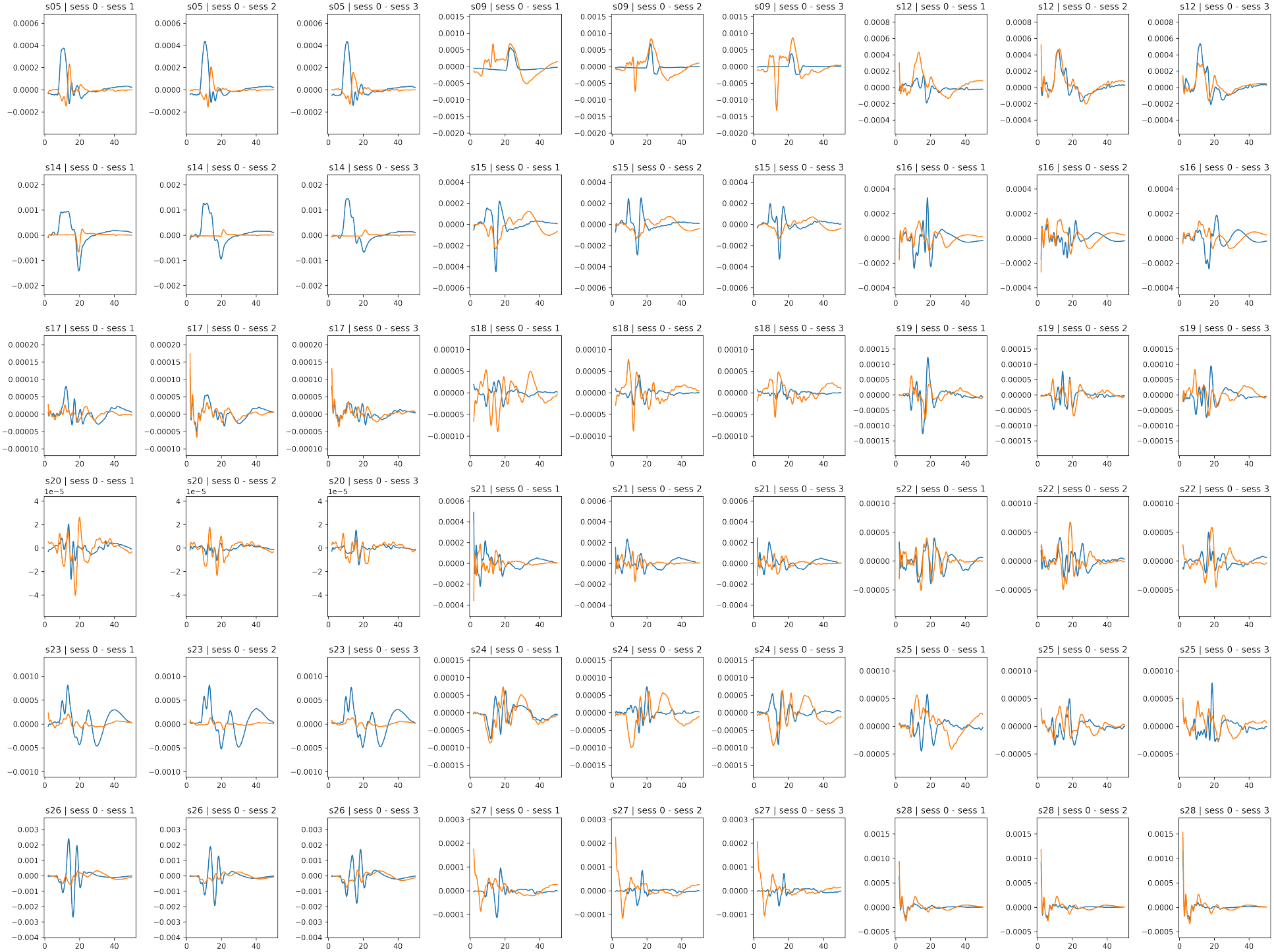
Within-participant differences between mean EMG waveforms in session 0 and each post-stimulation session. Each set of three adjacent panels corresponds to one participant and shows session 0 minus session 1, session 0 minus session 2, and session 0 minus session 3, respectively. Blue traces represent FCR and orange traces represent ECR; the horizontal axis indicates time relative to stimulus onset (ms), and the vertical axis indicates the difference in EMG amplitude (V). Difference waveforms were heterogeneous across participants, sessions, and muscles, with no uniform direction of change.

## Notes

### Competing Interest Statement

The authors have declared no competing interest.

### Funding

- MSU Institute for Artificial Intelligence

